# Meta-analysis of Public RNA-sequencing Data of Drought and Salt Stresses in Different Phenotypes of *Oryza sativa*

**DOI:** 10.1101/2024.08.06.605779

**Authors:** Mitsuo Shintani, Hidemasa Bono

**Affiliations:** Graduate School of Integrated Sciences for Life, Hiroshima University, Higashi-Hiroshima, Japan; Genome Editing Innovation Center, Hiroshima University, Higashi-Hiroshima, Japan

**Keywords:** *Oryza sativa*, Salt stress, Drought stress, RNA-sequencing, Meta-analysis

## Abstract

Environmental stresses, such as drought and salt, adversely affect plant growth and crop productivity. While many studies have focused on established components of stress signaling pathways, research on unknown elements remains limited. In this study, we collected RNA sequencing (RNA-Seq) data from *Oryza sativa* registered in public databases and conducted a meta-analysis integrating multiple studies. We analyzed 105 paired RNA-Seq datasets from resistant or susceptible *Oryza sativa* cultivars under salt and drought conditions to identify novel stress-responsive genes with common expression changes across these datasets. A meta-analysis identified 10 genes specifically upregulated in resistant cultivars and 12 specifically upregulated in susceptible cultivars under both drought and salt conditions. Furthermore, by comparing previously identified stress-responsive genes in *Arabidopsis thaliana*, we explored genes potentially involved in stress response mechanisms that are conserved across plant species. The genes identified in this data-driven study may serve as targets for future research and genome editing.

## 1. Introduction

Rice (*Oryza sativa*) is an important food in East Asia, South Asia, the Middle East, Latin America, and the West Indies, and it is estimated to account for more than one-fifth of the calories consumed by humans worldwide (Sharif et al., 2014). Rice is a crucial crop because it is high in calories and contains more abundant amounts of essential vitamins and minerals than other grains (Mohidem et al., 2022). However, rice production faces salinity and drought stress, which hinders its cultivation.

Salinity and drought are major abiotic stresses that inhibit plant growth and productivity (Angon et al., 2022; Billah et al., 2021; Kamran et al., 2019; Ma et al., 2020; Yang & Guo, 2018). Rice is susceptible to the negative effects of drought and salt damage, which lead to serious issues of reduced yields. Twenty percent of agricultural land currently used worldwide is affected by salt stress, and this percentage increases every year owing to anthropogenic and natural factors (C. Liu et al., 2022). High Na+ levels induce K+ and Ca2+ efflux from the cytoplasm, causing intracellular homeostatic imbalance, nutrient deficiency, oxidative stress, growth retardation, and cell death (Ahanger & Agarwal, 2017). In addition, high salt concentrations reduce photosynthesis through stomatal limitations, such as stomatal closure (Munns & Tester, 2008). Salt stress also has a negative effect on photosynthesis because of non-stomatal limitations, such as chlorophyll dysfunction, deficiency of photosynthetic enzyme proteins and membranes, and disruption of the ultrastructure of the chloroplast (Gengmao et al., 2015; Jiang et al., 2012; Mittal et al., 2012).

Drought represents a state of soil water deficit in which insufficient water is available for plants to fully grow and complete their life cycles. As soil water evaporates, salinity is concentrated in the soil solution, causing drought and salinity simultaneously (“Impacts of soil salinity/sodicity on soil-water relations and plant growth in dry land areas: A review,” 2021).

Across Asia, 20% of all rice-producing areas are affected by drought each year (Gowda et al., 2011). Several studies focusing on plant drought have shown that drought delays the flowering time of rice plants and consequently suppresses plant growth, resulting in reduced ear number and fertile fruit number, ultimately reducing yield (Ekanayake et al., 1989; G. Liu et al., 2010; Yue et al., 2005; Zou et al., 2007). Drought stress causes changes in the anti-oxidant and osmotic regulatory systems of rice, leading to the accumulation of antioxidants and osmotic regulators (Ouyang et al., 2010; X. Wang et al., 2019).

Previous studies identified several genes that contribute to salinity and drought stress in rice (Hassan et al., 2023; C. Liu et al., 2022; Panda et al., 2021). For example, the transcription factors *OsMYBc, OsbZIP71*, and *OsNF-YC13* improve salt tolerance by activating K+/Na+ transporters and increasing the K+/Na+ ratio (C. Liu et al., 2014; Manimaran et al., 2017; R. Wang et al., 2015).

In addition, *OsP5CS1* and *OsP5CR* play crucial roles in synthesizing compatible solute proline, increasing proline levels, and thereby enhancing salt tolerance (Sripinyowanich et al., 2013). *OsCPK4* and *OsCPK12*, which encode calcium-dependent protein kinases, are involved in scavenging reactive oxygen species (ROS), thereby enhancing salt tolerance (Asano et al., 2012; Campo et al., 2014). Similarly, the transcription factors *OsZFP179* and *OsZFP182* also improve salt tolerance by enhancing ROS scavenging ability (Huang et al., 2012; S.-J. Sun et al., 2010).

Regarding drought tolerance, the expression of *OsCPK9* promotes stomatal closure and improves osmotic regulation, thereby enhancing plant drought tolerance (Wei et al., 2017). The transcription factors *OsNAC5* and *OsNAC10* increase grain yield and root development under drought conditions (Jeong et al., 2010, 2013). *OsDREB2A* enhances the survival of transgenic plants under drought stress and *OsDREB2B* increases root length and number (Cui et al., 2011; Matsukura et al., 2010).

However, current research has only revealed a small part of the overall stress response mechanism in rice. In particular, the causes of large differences in stress sensitivity among rice varieties are not fully understood. Therefore, identifying the factors that determine the differences in stress sensitivity is important for elucidating the entire stress response mechanism in rice.

In this study, we focused on the differences in drought and salt stress responses among rice varieties and attempted to identify genes specifically involved in stress sensitivity (susceptible), stress tolerance (resistant), and each phenotype. Through the integrated analysis of a large amount of RNA sequencing (RNA-Seq) data retrieved from public databases, we explored novel responsive genes that have not been reported to be associated with salt or drought stress. By comparing differentially expressed genes identified between cultivars with different susceptibilities to stress, we narrowed down candidate genes involved in the phenotype.

In a previous study, we performed a meta-analysis of stress conditions involving abscisic acid (ABA), salt, and dehydration in *Arabidopsis thaliana* (Shintani et al., 2024). By comparing the stress-responsive genes identified in *Oryza sativa* from this study with those from the meta-analysis of *Arabidopsis thaliana*, we identified candidate genes that contribute to stress response mechanisms conserved across plant species.

The findings of this study provide a new perspective for basic research on the stress response mechanisms in rice. Moreover, the identified genes have the potential for use as target candidates for genome editing to develop stress-resistant rice.

## 2. Materials and methods

### 2.1 Manual curation of gene expression data from public databases

RNA-Seq data were collected from the National Center for Biotechnology Information Gene Expression Omnibus (NCBI GEO) (https://www.ncbi.nlm.nih.gov/geo/) (Barrett et al., 2013) and European Bioinformatics Institute BioStudies (EBI BioStudies) (https://www.ebi.ac.uk/biostudies/) (Sarkans et al., 2018). In the NCBI GEO, a comprehensive search was performed using the following query: (“drought”[All Fields] OR “water deficit”[All Fields] OR “salt”[All Fields] OR “NaCl”[All Fields] OR “salinity”[All Fields]) AND “*Oryza sativa*”[porgn] AND “Expression profiling by high throughput sequencing”[Filter].

In the EBI BioStudies, projects related to stress treatments were selected by searching with the following parameters: “Keywords: ‘RNA-Seq’ AND ‘*Oryza sativa*’ AND (‘salt’ OR ‘drought’),” “Collection: ‘Array Express’.”

Manual curation was performed based on the search results. The manual curation process included collecting metadata from the database, such as stress treatment conditions and sample tissues, along with the raw RNA-Seq data. Based on the descriptions in the papers and database metadata, the phenotype of each cultivar was classified as either “resistant” or “susceptible” to stress. Another criterion was the presence of paired stress-treated and stress-untreated control samples in the same project. This process resulted in the creation of 105 paired datasets from 202 samples from 11 projects.

### 2.2 Gene expression quantification

FASTQ-formatted files for each RNA-Seq run accession number were obtained using the prefetch and fastq-dump commands of the SRA Toolkit (v3.0.0) [https://github.com/ncbi/sra-tools]. Quality control of the raw reads was performed using Trim Galore (v0.6.7) [https://github.com/FelixKrueger/TrimGalore]. Transcript quantification was performed using Salmon (v1.8.0) [https://github.com/COMBINE-lab/salmon] (Patro et al., 2017), and the cDNA sequences downloaded from Ensembl Plants (release 56) (Oryza_sativa.IRGSP-1.0. cdna.all.fa) were used as reference. To ensure data consistency and enable accurate gene expression comparisons, we used the *Oryza sativa japonica* reference transcriptome for expression quantification in both *japonica* and *indica* datasets. This approach enhances comparability and unifies annotations. We chose the *japonica* reference because of its more comprehensive and well-curated gene annotations, which provided a rich foundation for functional analysis across both subspecies. As a result, the quantitative RNA-Seq data were calculated as the TPM. Transcript-level TPM values were summarized at the gene level using tximport (v1.26.1) [https://bioconductor.org/packages/release/bioc/html/tximport.html].

### 2.3 TN ratio and score calculation

Gene expression data from different experiments were normalized by calculating the TN ratio, which represents the ratio of gene expression between stress-treated (T) and non-treated (N) samples. If the TN ratio was higher than the threshold, the gene was considered to be upregulated; whereas if it was less than the reciprocal of the threshold, the gene was considered downregulated. Otherwise, the gene was considered unchanged. To classify the upregulated and downregulated genes, 1.5-fold, 2-fold, 5-fold, and 10-fold thresholds were evaluated, and a 2-fold threshold was adopted. Therefore, genes with a TN ratio higher than 2 were classified as upregulated, while genes with a TN ratio lower than 0.5 were classified as downregulated. The TN score of each gene was determined by subtracting the number of downregulated experiments from the number of upregulated experiments to assess changes in gene expression under stress conditions across multiple experiments. The TN ratio and TN score were computed using code that had been utilized in a previous study (Ono & Bono, 2021).

### 2.4 Gene set enrichment analysis

Gene set enrichment analysis was performed using the web tool ShinyGO (v0.77) [http://bioinformatics.sdstate.edu/go77/] (Ge et al., 2020) to analyze the DEG sets. For the *Oryza sativa* analysis, the species was set to “*Oryza sativa Japonica* Group,” and the pathway database was set to “GO Biological Process.” All other parameters were set to the default settings.

### 2.5 Visualization

A web-based Venn diagram tool [https://bioinformatics.psb.ugent.be/webtools/Venn/] and the UpSet plot tool [https://asntech.shinyapps.io/intervene/] (A. Khan & Mathelier, 2017) were used to search for and visualize overlapping genes.

### 2.6 Functional annotation of *Oryza sativa* genes

To functionally annotate the *Oryza sativa* genes, we retrieved the “Gene ID,” “Gene name,” and “Gene description” for each gene from the Ensembl Plants (release 56, *Oryza sativa Japonica* Group genes (IRGSP-1.0)). We then identified the putative orthologs of these genes in *Arabidopsis thaliana* by performing a BLASTP (v2.12.0) search using the *Oryza sativa* protein sequences (*Oryza_sativa*.IRGSP-1.0.pep.all.fa.gz) as queries against the *Arabidopsis thaliana* protein database obtained from Ensembl Plants (release 56, *Arabidopsis_thaliana*.TAIR10.pep.all.fa.gz). The BLASTP search was conducted with an E-value cutoff of 1e-10. For each *Oryza sativa* gene, the top hit in the *Arabidopsis thaliana* protein database was considered the putative ortholog. The corresponding “Gene ID,” “Gene name,” and “Gene description” for the *Arabidopsis thaliana* orthologs were obtained from the Ensembl Plants (release 56, *Arabidopsis thaliana* genes (TAIR10)).

## 3. Results

### 3.1 Overview of this study

Initially, we present an overview of the entire study. Through comprehensive data analysis, this research aimed to achieve two primary objectives:

1. Identification of genes suggested to contribute to stress tolerance phenotypes in *Oryza sativa*.
2. Identification of genes implicated by the data to be involved in common stress response mechanisms across plant species, specifically in *Oryza sativa* and *Arabidopsis thaliana*.

This study consisted of six steps as shown in Figure 1.

**Figure 1.**
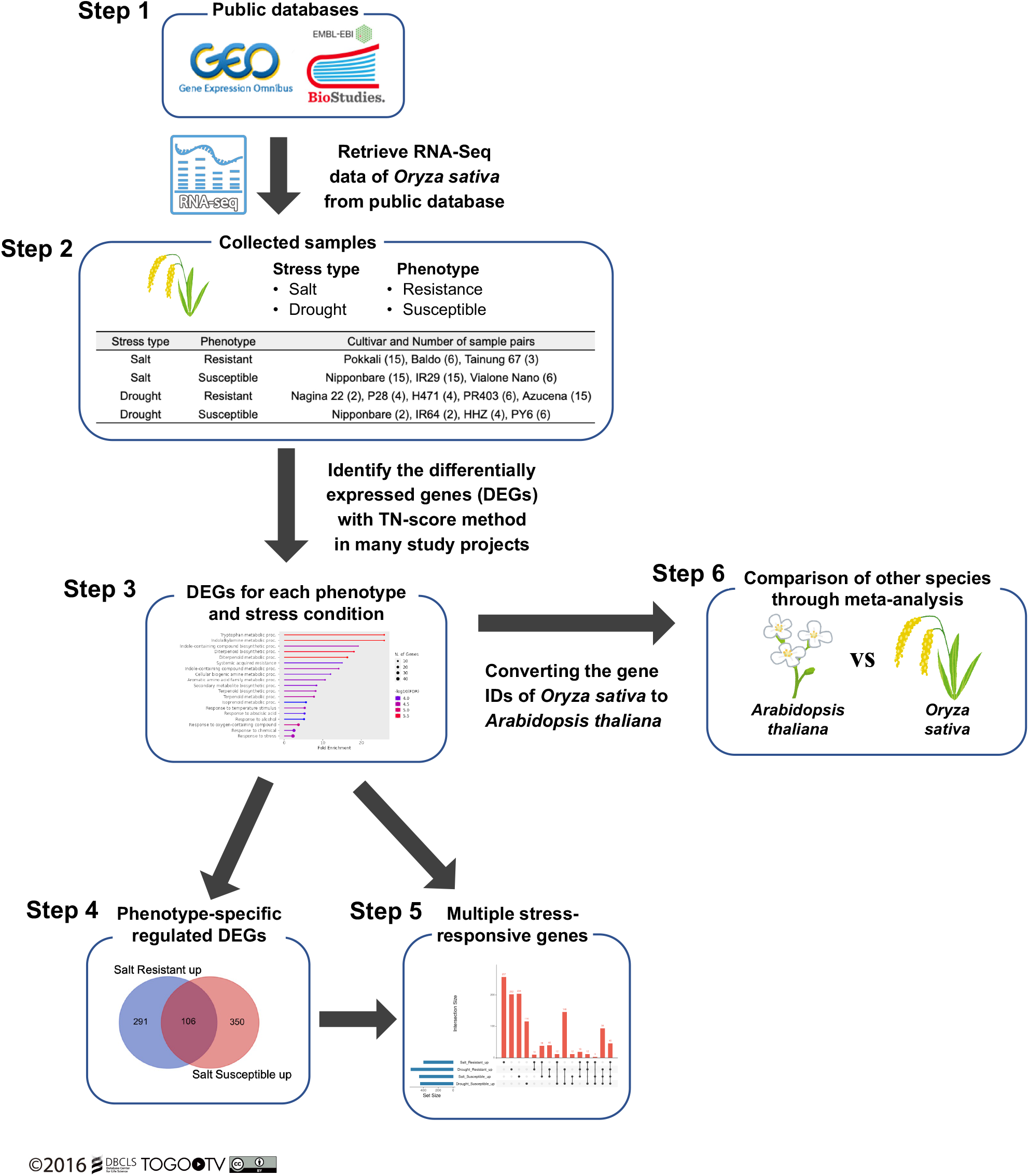
Overview of the six steps of this study. Step 1: RNA-Seq data for *Oryza sativa* were retrieved from public databases (National Center for Biotechnology Information Gene Expression Omnibus (NCBI GEO) and European Bioinformatics Institute BioStudies (EBI BioStudies). Step 2: Samples were categorized according to stress type (salt/drought) and phenotype (resistant/susceptible). Differentially expressed genes (DEGs) were identified using the TN-score method. Step 3: Enrichment analysis was performed on the DEGs for each phenotype and stress condition to evaluate the validity of the analysis. Step 4: Genes involved in the phenotypic differences between *Oryza sativa* cultivars were selected under drought and salt stress conditions. Step 5: Common genes among different stress conditions were selected. Step 6: Genes involved in stress response mechanisms conserved across different plant species were selected.

### 3.2 Retrieval and curation of RNA-Seq data from public databases

RNA-Seq data were retrieved from the National Center for Biotechnology Information Gene Expression Omnibus (NCBI GEO) and European Bioinformatics Institute BioStudies (EBI BioStudies) public databases. Compared with microarray data, RNA-Seq data were primarily derived from the Illumina platform, making them more suitable for comparative analyses of studies from different research groups. Therefore, only RNA-Seq data were used in this study, while microarray data were excluded. In this study, based on the description in the papers and metadata of databases, the phenotype of cultivars reported to present strong resistance was defined as “resistant” and the cultivars reported to present relatively weak resistance were defined as “susceptible.” A total of 202 samples were collected from the 11 projects, resulting in 105 paired datasets of stress-treated and untreated control samples. Because of manual curation, the collected data were divided into four categories based on the stress type and phenotype: (1) salt-resistant, (2) salt-susceptible, (3) drought-resistant, and (4) drought-susceptible. The number of paired datasets for each category was as follows: 24 for salt-resistant, 36 for salt-susceptible, 31 for drought-resistant, and 14 for drought-susceptible. To systematically organize the data used in the analysis, a table including the stress type, phenotype, subspecies, research project ID, and experimental pair number was prepared for each rice variety (Supplementary Table S1). When the number of collected sample pairs was compared among the tissues, it was revealed that samples derived from “leaf” contained the most data (Supplementary Figure S1). Details on all other metadata are presented in Supplementary Table S2.

### 3.3 Analysis of collected data and selection of differentially expressed genes in *Oryza sativa*

The collected samples were subjected to quality control and expression quantification using Salmon(Patro et al., 2017), which calculated the transcripts per million (TPM) values for each gene in each sample (Supplementary Table S3). Stress-treated and non-treated samples were paired for each gene, and the expression ratios (TN ratios) and TN scores were calculated. The TN ratio was calculated as follows:

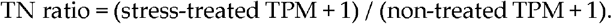

If the TN ratio was higher than the threshold, the gene was considered upregulated; whereas if it was lower than the reciprocal of the threshold, it was considered downregulated. Otherwise, it was considered unchanged. To classify the upregulated and downregulated genes, we evaluated 1.5-fold, 2-fold, 5-fold, and 10-fold thresholds and finally chose the 2-fold threshold. Therefore, genes with a TN ratio higher than 2 were classified as upregulated, while genes with a TN ratio lower than 0.5 were classified as downregulated. The TN score of each gene was determined by subtracting the number of downregulated experiments from the number of upregulated experiments to assess changes in gene expression under stress conditions across the entire experiment.

The collected samples were classified into the four aforementioned categories based on the phenotype and type of stress treatment, and the TN scores for each category were calculated. After considering multiple expression ratio thresholds, we adopted a 2-fold threshold (TN2). This threshold was slightly lower to provide a comprehensive analysis. More severe scores for the 5-fold (TN5) and 10-fold (TN10) thresholds were also calculated and are listed in the Supplementary Table. The TN ratios and TN scores for all the genes are available online (Supplementary Tables S4 and S5).

Additionally, the distribution of TN scores for the genes was visualized using scatter plots (Supplementary Figure S2). Focusing on points where scores changed significantly on the scatter plot, the top and bottom genes based on TN2 scores were selected as differentially expressed genes (DEGs). This accounted for the top or bottom 1% of all reference genes. Abrupt score changes indicated that the expression of these genes was remarkably variable in the dataset used in this study. The number of identified genes and range of TN2 scores are summarized in Table 1. Lists of the upregulated and downregulated genes are available online (Supplementary Tables S6 and S7, respectively). For the DEGs identified in rice, the corresponding orthologs in *Arabidopsis thaliana* were determined using sequence similarity searches with BLASTP. The list of these genes is available online (Supplementary Table S8).

**Table 1.**
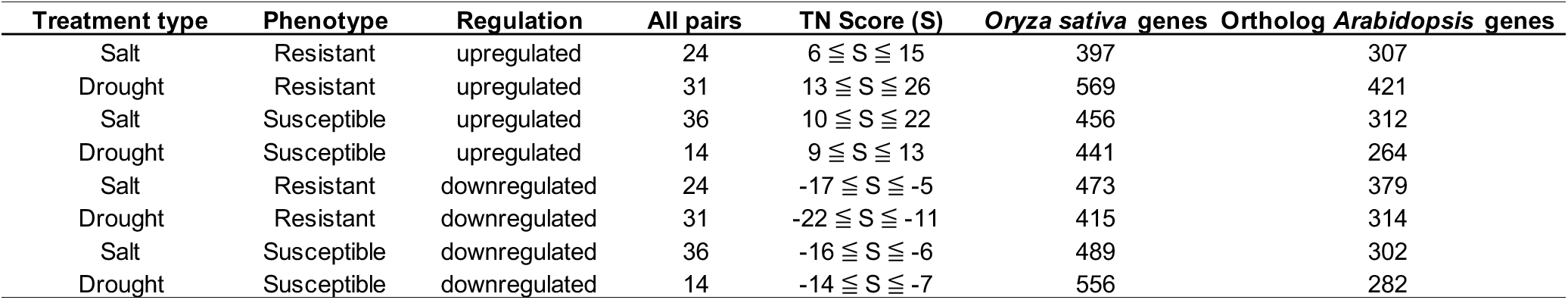
Distribution of DEG numbers and TN2 scores in *Oryza sativa* under different stress conditions and phenotypes abased on the meta-analysis.

### 3.4 Enrichment analysis to evaluate the characteristics of DEGs

To evaluate the characteristics of the identified DEGs, we performed enrichment analyses of individual DEG groups using ShinyGO. The results are shown in Supplementary Figure S3 and Supplementary Table S9. The top two GO terms for each DEG are listed in Supplementary Table S10.

Enrichment analysis revealed distinct patterns of gene regulation under different stress conditions and phenotypes. These results suggest that diverse stress response mechanisms may exist depending on the type of stress and phenotypic traits.

Furthermore, we focused on DEGs that showed similar expression patterns (upregulation or downregulation) across different phenotypes and stress conditions and considered them as a single group. We performed enrichment analyses for two types of combined DEGs. The first group consisted of all genes whose expression was upregulated under at least one condition (Supplementary Figure S4a), and the second group consisted of all genes whose expression was downregulated under at least one condition (Supplementary Figure S4b).

For upregulated DEGs (Supplementary Figure S4a), the most significantly enriched terms were “diterpenoid metabolic processes” and “water deprivation.” Other enriched terms included responses to various abiotic stressors, such as heat and salt stress, and hormone responses, especially abscisic acid (ABA). These results support the validity of the selection of DEGs responsive to salt and drought stress by the meta-analysis.

For the downregulated genes (Supplementary Figure S4b), the analysis revealed a different set of enriched terms. The most significantly enriched terms were related to nitrogen metabolism, including “nitrate metabolic processes,” “nitrate assimilation,” and “reactive nitrogen species metabolic processes.” Interestingly, several terms involved in oxidative stress response and detoxification were also significantly enriched, such as “hydrogen peroxide catabolic process” and “reactive oxygen species metabolic process.” Additionally, photosynthesis-related processes and transmembrane transport were among the enriched pathways for downregulated genes. Detailed results of the enrichment analysis for individual DEGs and all combined DEGs are provided in Supplementary Table S9.

### 3.5 Selection of genes involved in phenotypic differences between *Oryza sativa* cultivars under drought and salt stress conditions

We compared the DEGs between the stress-resistant and stress-susceptible phenotypes for both salt and drought stress to identify genes specific to each condition (Supplementary Figure S5). The numbers of common and unique genes in each group are summarized in Supplementary Table S11.

We conducted an enrichment analysis of these individual gene sets, and the results are shown in Supplementary Figure S5. However, as no hits were found in gene set (j), only those for the “All available gene sets” setting were used. The top GO terms for each DEG are summarized in Supplementary Table S12.

Interestingly, the enrichment analysis showed that the most significantly enriched term for both the upregulated genes in the salt susceptible and salt resistant groups as well as the drought susceptible and drought resistant groups was “GO:0009631 Cold acclimation.” This suggests that the importance of stress mechanisms of cold acclimation may be similar to that of both salinity and drought stress, regardless of the *Oryza sativa* phenotype.

In addition, in Supplementary Figure S5a, the most significantly enriched term in the enrichment analysis of the upregulated combine DEGs was “diterpenoid metabolic processes.” In contrast, the most significantly enriched term in the genes specifically downregulated in the drought-resistant phenotype was “diterpenoid biosynthesis (KEGG: osa00904).” This suggests that the pathways involving diterpenoids may regulate stress responses by upregulating or downregulating the expression of related genes. All gene lists, Venn diagrams visualizing overlapping genes, and enrichment results are shown in Supplementary Tables (Supplementary Table S13 and S14).

### 3.6 Selection of genes common among different stress conditions

The overlap of genes that were commonly upregulated or downregulated across the four categories based on the two stress conditions and two phenotypes each was visualized using UpSet plots (Figure 2).

**Figure 2.**
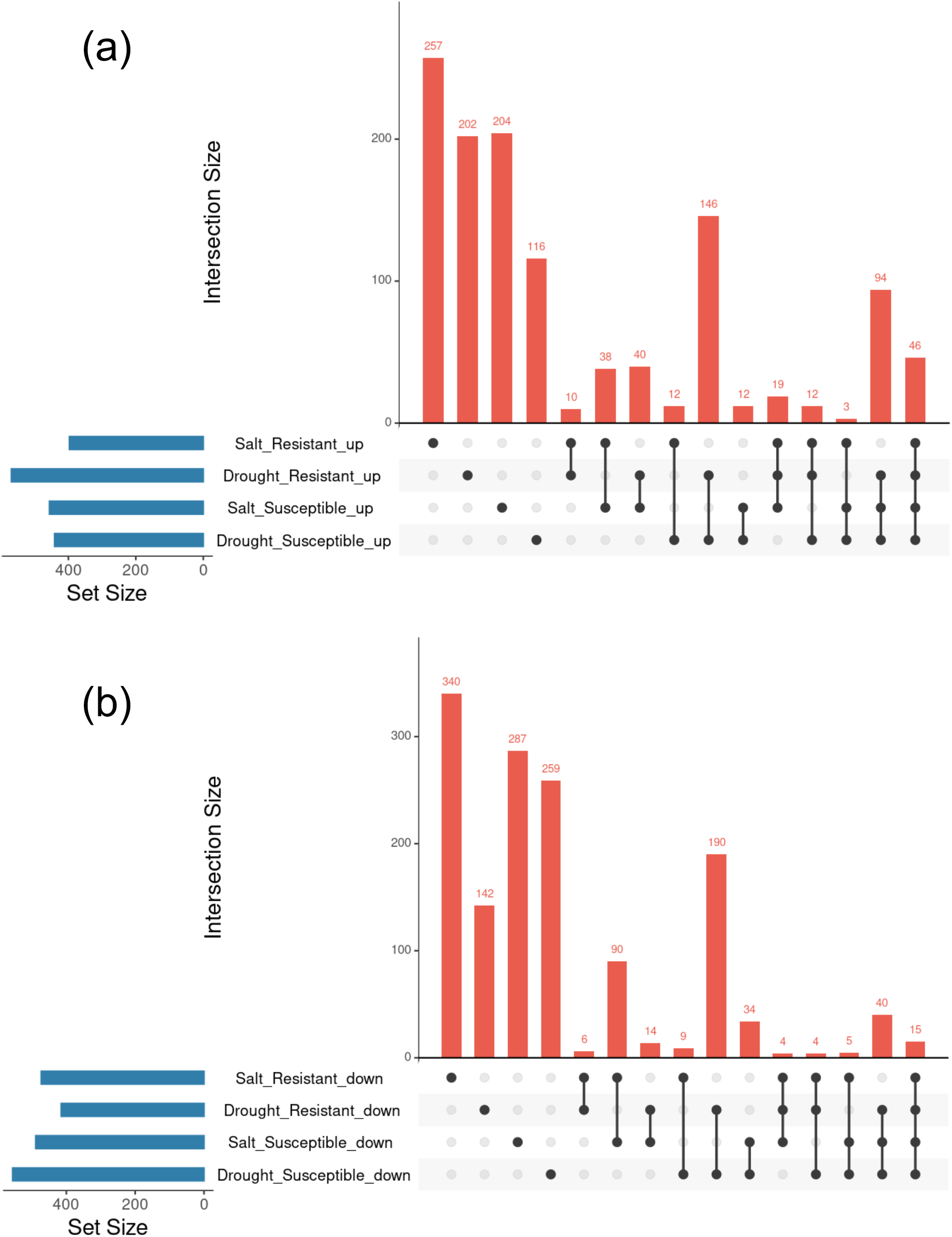
UpSet plots showing the overlap and specificity of DEGs in *Oryza sativa* under various stress conditions and phenotypes. UpSet plots illustrating the overlap of commonly or specifically regulated genes across the four categories combining salt and drought stress conditions with resistant and susceptible phenotypes. (a) Upregulated genes; (b) downregulated genes

All overlapping genes are listed in Supplementary Table 15. The number of genes whose expression specifically increased or decreased in each phenotype under both stress conditions is summarized in Supplementary Table S16.

Among these genes, those whose expression were upregulated in the resistant and susceptible varieties are summarized in Table 2.

**Table 2.** List of *Oryza sativa* genes that were specifically upregulated in the resistant or susceptible phenotype under both salt and drought stress conditions.

| Overlap | Total number of genes | Os-Gene stable ID | Os-Gene name | Os-Gene description |
| --- | --- | --- | --- | --- |
| Drought_Resistant_up Salt_Resistant_up | 10 | *Os12g0594000* | *OsWD40-198* | WD40 repeat-like domain containing protein |
|  |  | *Os09g0379600* | *Oshox25* | Similar to Homeobox-leucine zipper protein HOX25 |
|  |  | *Os03g0709300* | *OsUCL9* | Similar to Chemocyanin precursor (Basic blue protein) (Plantacyanin) |
|  |  | *Os02g0140400* | *OsOSC4* | Similar to Beta-amyrin synthase |
|  |  | *Os02g0139000* | *PHR3* | Transcription factor, Regulation of Pi signaling and homeostasis, Tolerance to low-Pi stress |
|  |  | *Os08g0290700* | - | Winged helix repressor DNA-binding domain containing protein |
|  |  | *Os11g0112400* | *prx130* | Peroxidase (EC 1.11.1.7) |
|  |  | *Os01g0767600* | - | Conserved hypothetical protein |
|  |  | *Os09g0555500* | *OsPSY3* | Similar to Chloroplast phytoene synthase 3 |
|  |  | *Os01g0155800* | - | Conserved hypothetical protein |
| Drought_Susceptible_up Salt_Susceptible_up | 12 | *Os01g0159800* | *OsbHLH035* | Similar to DNA binding protein |
|  |  | *Os05g0119100* | - | Hypothetical protein |
|  |  | *Os02g0209400* | - | Conserved hypothetical protein |
|  |  | *Os10g0450900* | *GRP* | Similar to Glycine-rich cell wall structural protein 2 precursor |
|  |  | *Os10g0517500* | *RRJ1* | Cys/Met metabolism, pyridoxal phosphate-dependent enzyme domain containing protein |
|  |  | *Os12g0510750* | - | Conserved hypothetical protein |
|  |  | *Os09g0278000* | - | Hypothetical conserved gene |
|  |  | *Os07g0673900* | *HIGD2* | Hypoxia induced protein, Early stage of hypoxia signaling |
|  |  | *Os11g0533400* | - | Conserved hypothetical protein |
|  |  | *Os11g0700900* | *C10923* | Glycoside hydrolase, subgroup, catalytic core domain containing protein |
|  |  | *Os03g0305100* | - | Similar to AMP-binding protein |
|  |  | *Os03g0184550* | - | Similar to Dihydroflavonol-4-reductase |

In total, 10 genes were consistently upregulated in the resistant phenotype under both stress conditions. These included genes involved in various cellular processes, such as transcription regulation (*Oshox25* and *PHR3*), secondary metabolism (*OsOSC4* and *OsPSY*), and peroxidase (*prx130*).

In contrast, 12 genes were consistently upregulated in the susceptible phenotype under both salt and drought stress conditions. These genes have different functional profiles, including a bHLH transcription factor (*OsbHLH035*), glycine-rich cell wall structural protein (*GRP*), hypoxia-induced gene (*HIGD2*), and enzymes involved in various metabolic processes (*RRJ1* and *C10923*).

### 3.7 Selection of genes involved in stress response mechanisms conserved across different plant species

In a previous study, we performed a meta-analysis of *Arabidopsis thaliana* under ABA, salt, and dehydration conditions and identified genes that were upregulated or downregulated under each condition (Shintani et al., 2024). Here, we used *Arabidopsis thaliana* orthologous genes corresponding to the *Oryza sativa* genes identified in this study to determine whether any genes shared common. The overlap in expression of commonly regulated genes was visualized using UpSet plots (Supplementary Figure S6). Satisfying the above criteria, 11 genes were upregulated (At_ABA_up, At_Dehydration_up, At_Salt_up, Os_Drought_Resistant_up, Os_Drought_Susceptible_up, Os_Salt_Resistant_up, and Os_Salt_Susceptible_up) and 1 gene was downregulated (At_ABA_down, At_Dehydration_down, At_Salt_down, Os_Drought_Resistant_down, Os_Drought_Susceptible_down, Os_Salt_Resistant_down, and Os_Salt_Susceptible_down) under salt and drought conditions (Table 3). These genes may be involved in the stress response mechanisms that are common across different plant species.

**Table 3.** Comparison of ABA, salt, and drought stress-responsive genes conserved in rice and *Arabidopsis*.

| Overlap | Total number<br>of genes | At-Gene<br>stable ID | At-Gene<br>Name | At-Description |
| --- | --- | --- | --- | --- |
| At_ABA_up At_Dehydration_up At_Salt_up Os_Drought_Resistant_up Os_Drought_Susceptible_up Os_Salt_Resistant_up Os_Salt_Susceptible_up | 11 | AT1G52690 | LEA7 | Late embryogenesis abundant protein (LEA) family protein |
|  |  | AT3G03341 | - | cold-regulated protein |
|  |  | AT1G07430 | HAI2 | highly ABA-induced PP2C protein 2 |
|  |  | AT2G35300 | LEA18 | Late embryogenesis abundant protein, group 1 protein |
|  |  | AT3G48510 | - | AtIII18x5-like protein |
|  |  | AT2G39050 | EULS3 | hydroxyproline-rich glycoprotein family protein |
|  |  | AT3G24520 | HSFC1 | heat shock transcription factor C1 |
|  |  | AT2G46270 | GBF3 | G-box binding factor 3 |
|  |  | AT5G66400 | RAB18 | Dehydrin family protein |
|  |  | AT2G47770 | TSPO | TSPO(outer membrane tryptophan-rich sensory protein)-like protein |
|  |  | AT2G46680 | HB-7 | homeobox 7 |
| At_ABA_down At_Dehydration_down At_Salt_down Os_Drought_Resistant_down<br>Os_Drought_Susceptible_down Os_Salt_Resistant_down Os_Salt_Susceptible_down | 1 | AT5G05440 | PYL5 | Polyketide cyclase/dehydrase and lipid transport superfamily protein |
| At_ABA_up At_Dehydration_up At_Salt_up Os_Drought_Resistant_up Os_Salt_Resistant_up | 6 | AT4G19230 | CYP707A1 | cytochrome P450, family 707, subfamily A, polypeptide 1 |
|  |  | AT1G60190 | PUB19 | ARM repeat superfamily protein |
|  |  | AT1G45249 | ABF2 | abscisic acid responsive elements-binding factor 2 |
|  |  | AT3G63060 | EDL3 | EID1-like 3 |
|  |  | AT1G29640 | - | senescence regulator (Protein of unknown function, DUF584) |
|  |  | AT1G48000 | MYB112 | myb domain protein 112 |
| At_ABA_down At_Dehydration_down At_Salt_down Os_Drought_Resistant_down Os_Salt_Resistant_down | 1 | AT1G80050 | APT2 | adenine phosphoribosyl transferase 2 |

For the *Oryza sativa* genes in the ‘resistant’ phenotype category used in this study, six genes were upregulated (At_ABA_up, At_Dehydration_up, At_Salt_up, Os_Drought_Resistant_up, and Os_Salt_Resistant_up) and one gene was downregulated (At_ABA_down, At_Dehydration_down, At_Salt_down, Os_Drought_Resistant_down, and Os_Salt_Resistant_down) was downregulated under salt and drought conditions (Table 3). These genes may be involved in stress responses that are common across plant species, particularly in mechanisms important for stress tolerance.

The list of *Oryza sativa* gene IDs identified in this study was converted to *Arabidopsis thaliana* IDs and compared with previous studies. All gene lists are available online (Supplementary Table S17).

## 4. Discussion

This study aimed to identify novel genes and potential pathways involved in drought and salt stress responses in *Oryza sativa* using a meta-analysis of publicly available RNA-Seq data. By integrating datasets from multiple studies covering different *Oryza sativa* cultivars and stress conditions, we aimed to reveal robust and consistent gene expression patterns associated with stress resistance or susceptibility. Analysis of the tissue types of the collected samples revealed that while salt and drought stress are expected to have significant effects on the roots, the majority of the affected samples in this study consisted of “leaf” and “shoot” tissues, with a limited number of “root-”specific samples. Although this study focused on phenotypes and stress types rather than tissue specificity, tissue-specific analyses will likely become possible in the future as the amount of publicly available RNA-Seq data increases.

This study focused on several *Oryza sativa* cultivars that exhibited different phenotypic responses to salt and drought stress. The phenotypic classifications of “resistant” and “susceptible” used in this study are based solely on existing research reports and may not fully capture the nuanced stress tolerance levels of each cultivar. In addition, differences in stress treatment methods among the studies should be considered.

To minimize the impact of these differences and provide a robust comparative analysis of multiple datasets, we adopted the following approach. First, we directly compared the gene expression ratios between stressed and control samples in the same study. Subsequently, we calculated the TN score, which facilitated integration and comparative analysis of the data across multiple studies.

Based on the TN score, DEGs were identified for each dataset and categorized by stress type and phenotype. Enrichment analysis of each DEG set revealed the distinct and characteristic terms associated with each DEG group. Furthermore, we performed an enrichment analysis of genes that were upregulated or downregulated, irrespective of the phenotypic classification or stress type. The most significantly enriched term for upregulated genes was “diterpenoid metabolic process,” with 13 genes included in this category (Figure 3(a)). Diterpenoid phytoalexins have been shown to be potentially involved in the resistance of cereal plants to environmental stresses, such as drought (Murphy & Zerbe, 2020). The genes are listed in Supplementary Table S18.

These genes included those whose functions in stress mechanisms have been elucidated in previous studies. *OsCPS4* encodes a key enzyme that contributes to the biosynthesis of labdane-related diterpenoid (LRD) phytoalexins in *Oryza sativa*. Mutant analysis showed that the loss of function of *OsCPS4* significantly increased sensitivity to drought stress, suggesting that LRD biosynthesis involving *OsCPS4* contributes to the regulation of stomatal closure, independent of the ABA pathway (J. Zhang et al., 2021). The genes with unknown functions listed in this study may include those that contribute to the stress response through their involvement in LRD metabolism. Interestingly, in addition to the upregulated DEGs, the term enriched in genes specifically downregulated in the drought stress susceptible samples was “diterpenoid biosynthesis (KEGG: osa00904)” (Supplementary Figure S5 (d), Supplementary Table S18). This contrasting expression pattern suggests a complex regulatory mechanism in the diterpenoid pathway. In other words, the genes in this pathway may show diverse expression patterns depending on the type of stress, plant phenotype, and cultivar.

We also focused on a group of genes whose expression was upregulated in both the resistant and susceptible cultivars under the two stresses. Some of the genes commonly upregulated under salt and drought stress in the resistant phenotype included *OsPSY3* (*Os09g0555500*). PSY is a rate-limiting enzyme in carotenoid synthesis and represents a precursor required for ABA synthesis in various plants (Zhou et al., 2022). *OsPSY3* has been demonstrated in previous studies using reverse transcription quantitative PCR (RT-qPCR) experiments to show upregulation of gene expression after dehydration, salt, and ABA treatment (Du et al., 2015; Welsch et al., 2008). Therefore, the selection results were consistent with those of existing studies. The identification of this gene as one of the stress-responsive genes in the previous studies provides the evidence supporting the results of the meta-analysis. Other upregulated genes included transcription factor *OsPHR3* (*Os02g0139000*). This gene is also upregulated in response to dehydration stress and confirmed by RT-qPCR in the previous study (Abdirad et al., 2022). The expression of this gene is also induced by inorganic phosphate (Pi) deficiency and plays a role in improving tolerance to Pi deficiency as well as regulating nitrogen (N) homeostasis (Guo et al., 2015; Ruan et al., 2017; Y. Sun et al., 2018). Deficiency of Pi or N induces various morphophysiological adaptive responses and gene expression regulation in plants (Bechtaoui et al., 2021; F. Khan et al., 2023; Sakuraba, 2022; Xing et al., 2023). Although several studies have suggested a relationship between Pi and N deficiency and the ABA pathway and its biosynthesis, many aspects remain unclear and further research is needed (Fang Zhu et al., 2018; Hsieh et al., 2018; Jaskolowski & Poirier, 2024; Zakari et al., 2020; Yu Zhang et al., 2022).

In contrast, genes specifically upregulated in susceptible cultivars included *OsbHLH035* (*Os01g0159800*) and *RRJ1* (*Os10g0517500*). A previous meta-analysis study using transcriptome data suggested that genes such as *OsbHLH035* and *RRJ1* were upregulated by multiple heavy metal stresses, including mercury (Hg) and lead (Pb), and were subsequently validated by RT-qPCR (Fan et al., 2021). Drought-induced reductions in soil moisture can increase the concentration of heavy metals in the soil, which can have a synergistic negative effect on plants (Islam & Sandhi, 2022). Therefore, focusing on genes such as *OsbHLH035* and *RRJ1* is likely to provide important clues for understanding the combined effects of heavy metal stress and drought stress and for developing resistant varieties. However, it should be noted that experimental verification of the expression of these genes under salt or drought treatment has not yet been conducted.

Finally, a comparison with previous *Arabidopsis* meta-analysis data revealed several genes whose expression was altered in *Oryza sativa* and *Arabidopsis thaliana* under both salt and drought stress conditions (Table 3). To focus on genes involved in mechanisms commonly conserved between plant species regardless of the stress phenotype, we focused on genes shared by seven DEGs, with three in *Arabidopsis thaliana* and four in *Oryza sativa*. For example, *HSFC1* (*AT3G24520*) represented a gene commonly upregulated in both species, regardless of phenotype.

Previous studies on *Arabidopsis thaliana* have suggested that the expression of HSFC1 is involved in cold responses via a pathway independent of the C-repeat binding factor (CBF), a well-known transcription factor that controls cold acclimation in plants (Jia et al., 2016; Zhao et al., 2016).

The ortholog of Arabidopsis HSFC1 gene in *Oryza sativa* was *HSfC2b* (*Os06g0553100*). However, the functions and roles of these genes under salinity and drought conditions remain unclear. Heat shock factors (HSFs) that contribute to environmental stress tolerance occur in both *Arabidopsis thaliana* and *Oryza sativa* (Wenjing et al., 2020; Yan Zhang et al., 2022). Given the consistent upregulation of *HSFC1* and its ortholog *HSfC2b* across different stress conditions and plant species, these genes are promising candidates for further investigation.

This study had some limitations. First, because the identification of DEGs was based on the selection of TN scores rather than statistical tests, the results should be interpreted with caution. Second, the RNA-Seq data used in this study were collected under different growth and experimental conditions. Third, experimental validation using plant samples exposed to specific stress environments was not performed, indicating that further research is required to elucidate the functions of these genes. These limitations should be considered when interpreting our results and planning future studies.

## 5. Conclusion

Through a meta-analysis of publicly available RNA-Seq data, we obtained a list of potential target genes related to drought and salt stress responses in *Oryza sativa* and identified candidate genes that may influence stress-related phenotypes. These findings provide a valuable resource for future studies aimed at understanding the molecular mechanisms underlying stress resistance in rice and developing new strategies to improve crop productivity under adverse environmental conditions.

## Supplementary Materials

The Supplementary Material for this article can be found online at https://doi.org/10.6084/m9.figshare.25547083.v2.

## Acknowledgments

Computations were performed at the Hiroshima University Genome Editing Innovation Center.

## Author Contributions

Conceptualization, M.S. and H.B.; Methodology, M.S. and H.B.; Software, M.S. and H.B.; Validation, M.S. and H.B.; Formal Analysis, M.S. and H.B.; Investigation, M.S.; Resources, M.S. and H.B.; Data Curation, M.S.; Writing — Original Draft Preparation, M.S.; Writing — Review and Editing, M.S. and H.B.; Visualization, M.S.; Supervision, H.B.; Project Administration, H.B.; Funding Acquisition, H.B. All authors have read and agreed to the published version of the manuscript.

## Data Availability Statement

The data presented in this study are publicly available at Figshare (https://doi.org/10.6084/m9.figshare.25547083.v2).

## Conflicts of Interest declarations in manuscripts

The authors declare that this study was conducted in the absence of any commercial or financial relationships that could be construed as potential conflicts of interest.

